# Comparative evaluation of bolus and fractionated administration modalities for two antibody-cytokine fusions in immunocompetent tumor-bearing mice

**DOI:** 10.1101/712158

**Authors:** Emanuele Puca, Roberto De Luca, Frauke Seehusen, Josep Maria Monné Rodriguez, Dario Neri

## Abstract

Antibody-cytokine fusion proteins are being considered as biopharmaceuticals for cancer immunotherapy. Tumor-homing cytokine fusions typically display an improved therapeutic activity compared to the corresponding unmodified cytokine products, but toxicity profiles at equivalent doses are similar, since side effects are mainly driven by the cytokine concentration in blood. In order to explore avenues to harness the therapeutic potential of antibody-cytokine fusions while decreasing potential toxicity, we compared bolus and fractionated administration modalities for two tumor-targeting antibody-cytokine fusion proteins based on human interleukin-2 (IL2) and murine tumor necrosis factor (TNF) (i.e., L19-hIL2 and L19-mTNF) in two murine immunocompetent mouse models of cancer (F9 and C51). A comparative quantitative biodistribution analysis with radio-labeled protein preparations revealed that a fractionated administration of L19-hIL2 could deliver comparable product doses to the tumor with decreased product concentration in blood and normal organs, compared to bolus injection. By contrast, L19-mTNF (a product that causes a selective vascular shutdown in the tumor) accumulated most efficiently after bolus injection. Fractionated schedules allowed the safe administration of a cumulative dose of L19-mTNF, which was 2.5-times higher than the lethal dose for bolus injection. Dose fractionation led to a prolonged tumor growth inhibition for F9 teratocarcinomas, but not for C51 colorectal tumors, which responded best to bolus injection. Thus, dose fractionation may have different outcomes for the same antibody-cytokine product in different biological contexts.

## Introduction

The successful treatment of a growing number of tumor types with immune checkpoint inhibitors [1–9] has stimulated the interest in the exploration of alternative and complementary approaches for the selective boosting of T cell and NK cell activity against cancer. For many years, pro-inflammatory cytokines have been considered as a promising class of biopharmaceuticals for immunotherapy applications, but the practical applicability of cytokine therapies has been limited by severe toxicity at low doses, preventing escalation to therapeutically active regimens [10,11]. In spite of these limitations, some products have gained marketing authorization and have provided a therapeutic benefit for a small proportion of patients. Recombinant IL2 (aldesleukin, Proleukin^®^) obtained marketing authorization for the treatment of metastatic melanoma and for renal cell carcinoma [12]. Treatment with aldesleukin induced a prolonged overall survival only in a relatively small proportion of cancer patients, however substantial toxicities (e.g. vascular leak syndrome, nausea and fever) are frequently observed [13–15]. Recombinant TNF (Beromun^®^) has been approved for the treatment of soft-tissue sarcoma patients only for isolated limb perfusion procedures [16–19], whereas recombinant interferons have received marketing authorization for the treatment of various types of malignancies [20,21].

A number of strategies have been considered for the increase of the therapeutic index of pro-inflammatory cytokine payloads, including modification with cleavable polymers (*de facto* generating pro-drugs which regain activity upon hydrolytic loss of the polymer chains) [22] and antibody-cytokine fusions [23,24]. We and others have previously shown that antibody-cytokine fusions, capable of selective localization at the tumor site, can substantially increase therapeutic activity compared to the non-modified cytokine counterparts, when administered at equimolar doses [25–27]. L19-hIL2 and L19-hTNF are two products, which target the alternatively-spliced EDB domain of fibronectin (a marker of tumor angiogenesis) and which are currently being investigated in Phase III clinical trials for the treatment of Stage III B,C melanoma (EudraCT number: 2015-002549-72 and NCT03567889) and of metastatic soft-tissue sarcoma (EudraCT number: 2016-003239-38 and NCT03420014) [28–33].

Interleukin-2 is a monomeric protein which can easily be fused to antibodies in diabody format, a homobivalent protein architecture that exhibits superior tumor homing properties compared to monomeric scFv fragments [34]. Tumor necrosis factor features a non-covalent homotrimeric structure. ScFv fragments are normally used for the production of TNF fusions, since the use of non-covalent oligomeric structures (e.g., diabodies) would lead to TNF-mediated polymerization. Fusions of TNF with scFv fragments lead to the formation of stable non-covalent homotrimers, which can display excellent tumor-targeting properties in biodistribution studies as a result of rapid extravasation and multivalent antigen recognition [25,35–37]. The biological features of IL2- and TNF-based products, in the context of cancer immunotherapy, are considerably different. IL2 boosts the number and activity of intratumoral NK and T cells [38], while TNF displays a more complex mechanism of action, which involves an increase in vascular permeability leading to haemorrhagic necrosis within the neoplastic mass [25,35–37].

While antibody-cytokine fusions can increase the therapeutic index of their payload, toxicities at equimolar doses are often comparable. This feature, which may initially appear as paradoxical, is easy to understand considering the fact that only a small amount of product (in absolute terms) reaches the tumor mass (approximately 5-20 percent injected dose per gram in mice and 0.01-0.1 percent injected dose per gram in patient) [39,40]. Exposure of blood and normal organs to pro-inflammatory cytokines may cause hypotension, flu-like symptoms, nausea and vomiting [13–21]. Intense research activities are devoted to the search for methods which preserve the therapeutic activity of engineered cytokine products, while decreasing systemic toxicity [24]. Experimental strategies may include the allosteric activation of cytokine activity upon antigen binding [41], the design of split-cytokine fusions [42], the generation of dual-cytokine fusions [36,43–48], the selective depotentiation of cytokines by mutagenesis [36,49–51] and the use of masking antibodies to cover portions of the cytokine surface [52].

The toxicity of cytokine therapeutics is often manifested only at concentrations above a certain critical threshold and follows a non-linear profile [22]. It has been argued that a transient inhibition of cytokine activity at early time points after intravenous administration may represent a viable strategy for the generation of better tolerated products [24]. Here, we describe an experimental approach for the administration of high doses of antibody-cytokine fusion proteins, which would be lethal upon bolus injection. We reasoned that a fractionated administration in mice (or a slow infusion in patients) rather than a bolus injection should deliver comparable amounts of cytokine activity to the tumor site and keep blood concentrations below a critical threshold. We used L19-hIL2 [26] and L19-mTNF (featuring a murine cytokine moiety) [25] since these two products act by different mechanisms of action and since their full-human counterparts are being investigated in pivotal clinical trials. Quantitative biodistribution studies with radio-iodinated protein preparations and therapy experiments in tumor-bearing mice provided insights on the usefulness of administering fractionated product doses.

## Materials and Methods

### Tumor cell lines and reagents

All cell lines were obtained from the ATCC with the exception of C51 colon carcinoma (kindly provided by M.P. Colombo, Istituto Nazionale Tumori, Milan, Italy). Cell lines were received between 2015 and 2017, expanded, and stored as cryopreserved aliquots in liquid nitrogen. Cells were grown according the supplier’s protocol and kept in culture for no longer than 14 passages. Authentication of the cell lines also including check of postfreeze viability, growth properties, and morphology, test for mycoplasma contamination, isoenzyme assay, and sterility test were performed by the cell bank before shipment.

The clinical stage L19-hIL2 was kindly provided by Philogen S.p.A (Siena, Italy). The production and purification of the murine surrogate L19-mTNF was performed as described before [25].

### Biochemical characterization

Purified proteins were analyzed by size-exclusion chromatography on a Superdex 200 increase 10/300 GL column on an ÄKTA FPLC (GE Healthcare, Amersham Biosciences). SDS-PAGE was performed with 10% gels (Invitrogen) under reducing and non-reducing conditions. ESI/LC-MS was performed with an ACQUITY UPLC H class system equipped with an ACQUITY BEH300 C4 column (2.1 50 mm, 1.7-m particle size) sequentially coupled to a Waters Xevo G2-XS QTof ESI mass analyzer. Aminoacid sequences of L19-hIL2 and L19-mTNF can be found in **Supplementary Figure 1**.

### Immunofluorescence studies

Antigen expression was confirmed on ice-cold acetone fixed 8 μm cryostat sections of C51 and F9 tumors stained with IgG (L19)-FITC (final concentration 2 μg/mL) and detected with a Rabbit anti-FITC (Bio-Rad; 4510-7804) and Donkey anti-Rabbit AlexaFluor 488 (Invitrogen; A11008). For vascular staining Goat anti-CD31 (R&D System; AF3628) and Donkey anti-Goat AlexaFluor 594 (Invitrogen + A21209) antibodies were used. Slides were mounted with fluorescent mounting medium and analysed with fluorescent mounting medium (Dako Agilent).

### Animal and tumor models

Six to eight-week-old female BALB/c mice and 129/SvEv mice were obtained from Janvier Labs. Tumor cells were implanted subcutaneously in the flank using 1.5×10^7^ cells (F9), 1×10^6^ cells (C51). Experiments were performed under a project license (license number 04/2018) granted by the Veterina◻ramt des Kantons Zu◻rich, Switzerland, in compliance with the Swiss Animal Protection Act (TSchG) and the Swiss Animal Protection Ordinance (TSchV).

### Biodistribution studies

The capability of targeting EDB *in vivo* was assessed by quantitative biodistribution analysis. Purified protein samples (50-100 μg) were radio-iodinated with ^125^I (0.2 mCi per 100 μg of protein) and Chloramine T hydrate and purified on a PD10 column, according to a previously described procedure [53]. Radio-labeled proteins were injected into the lateral tail vein of 129/SvEv mice bearing subcutaneously implanted F9 lesions. Injected dose per mouse for L19-hIL2 were 30 μg and 10 μg for bolus and fractionated administration, respectively. Doses for L19-mTNF were 6 μg (bolus) and 3 μg (fractionated). For the bolus treatment group mice were sacrificed 24 hours after the bolus injection. For the fractionated treatment group mice were injected every 24 hours and sacrificed 24 hours after the last injection. Organ samples were weighed and radioactivity was counted using a gamma counter. The protein uptake in the different organs was calculated and expressed as the percentage of the injected dose per gram of tissue (%ID/g ± SEM n = 5 mice/group). Data were corrected for tumor growth and no radioactivity decay adjustments were made as the half-life of ^125^I is 60 days.

### Therapy studies

Mice were monitored daily and tumor volume was measured with a caliper (volume = length × width^2^ × 0.5). When tumors reached a suitable volume (approx. 70–100 mm^3^), mice were injected into the lateral tail vein with the pharmacological agents.

L19-hIL2 was administered at 100 μg and 400 μg once. L19-mTNF was dissolved in PBS, also used as negative control, and administered either as a single 10 μg bolus injection or as a 5 μg fractionated dose, which was repeated three times every 24 hours for a first experiment in F9 tumor bearing mice. In a second experiment in F9 tumors the single bolus injection was reduced at a dose of 5 μg, whereas the fractionated dose (5 μg) was extended to 5 cycles every 24 hours. In the C51 tumor model L19-mTNF was administered either as a single 3 μg bolus injection or as a 1 μg or 1.5 μg fractionated dose, which was repeated three times every 24 hours. Results are expressed as tumor volume in mm^3^ ± SEM and % mean body weight change ± SEM. For therapy experiments n = 4-5 mice/group.

### Plasma cytokine quantification

F9 tumor bearing mice were injected with either PBS, 5 μg L19-mTNF once or 3 μg L19-mTNF three times every 24 hours. Mice were sacrificed 24 hours after the last injection and blood was collected in Microtainers™ Tubes with Lithium Heparin (BD 365965). Tubes were centrifuged for 3 minutes at 2000g at 4°C. Plasma was collected and frozen at −80°C until cytokine level’s quantification was performed (Cytolab, Regensdorf, Switzerland).

### Histopathological evaluation of 129/SvEv mice

F9 teratocarcinoma bearing mice were euthanized 24 h after a single bolus injection of 400 μg L19-hIL2, 5 μg L19-mTNF or saline, and 24 h after the three daily injections of 200 μg L19-hIL2 or 3 μg L19-mTNF. A full necropsy was conducted in each mouse, main organs were removed and fixed in 4% neutral-buffered formalin (Formafix, Hittnau, Switzerland) for 48 hr. After fixation tissues were trimmed, dehydrated in graded alcohol and routinely paraffin wax embedded. Consecutive sections (3–5 μm thick) were prepared, mounted on glass slides and routinely stained with hematoxylin and eosin (HE).

## Results and Discussion

**Figure 1A** presents a schematic representation of possible patterns of fibronectin splice variant expression in subluminal aspects of tumor blood vessels. EDB-containing fibronectin can, in principle, be found in the basement membrane of tumor blood vessels and/or in the tumor interstitium and stroma. These two distinct features can indeed be found in patient specimens [54–57], as well as in F9 teratocarcinomas and C51 colorectal tumors: two murine models of cancer that were used in this study. A predominantly vascular pattern of EDB expression was observed in F9 tumors, while a diffuse staining of the interstitium was seen in C51 tumors [**Figure 1B**]. We used L19-hIL2 [26] and L19-mTNF [25] as representative immunostimulatory fusion proteins, in order to test the impact of dose fractionation in different models of cancer. **Figure 1C** **and** **D** show the molecular formats, SDS-PAGE and gel filtration profiles of L19-hIL2 and L19-mTNF, respectively. The non-covalent homodimer (L19-hIL2) and homotrimer (L19-mTNF) run as monomers in SDS-PAGE, as expected because of the denaturing conditions of this technique. The masses of the monomers were confirmed by Mass Spectrometry [**Supplementary Figure 3**], and the multimeric products showed in both cases a pure peak with the correct size (i.e., based on the elution volumes of standard proteins provided in the manufacturer’s brochure) in size exclusion chromatography.

**Figure 1:**
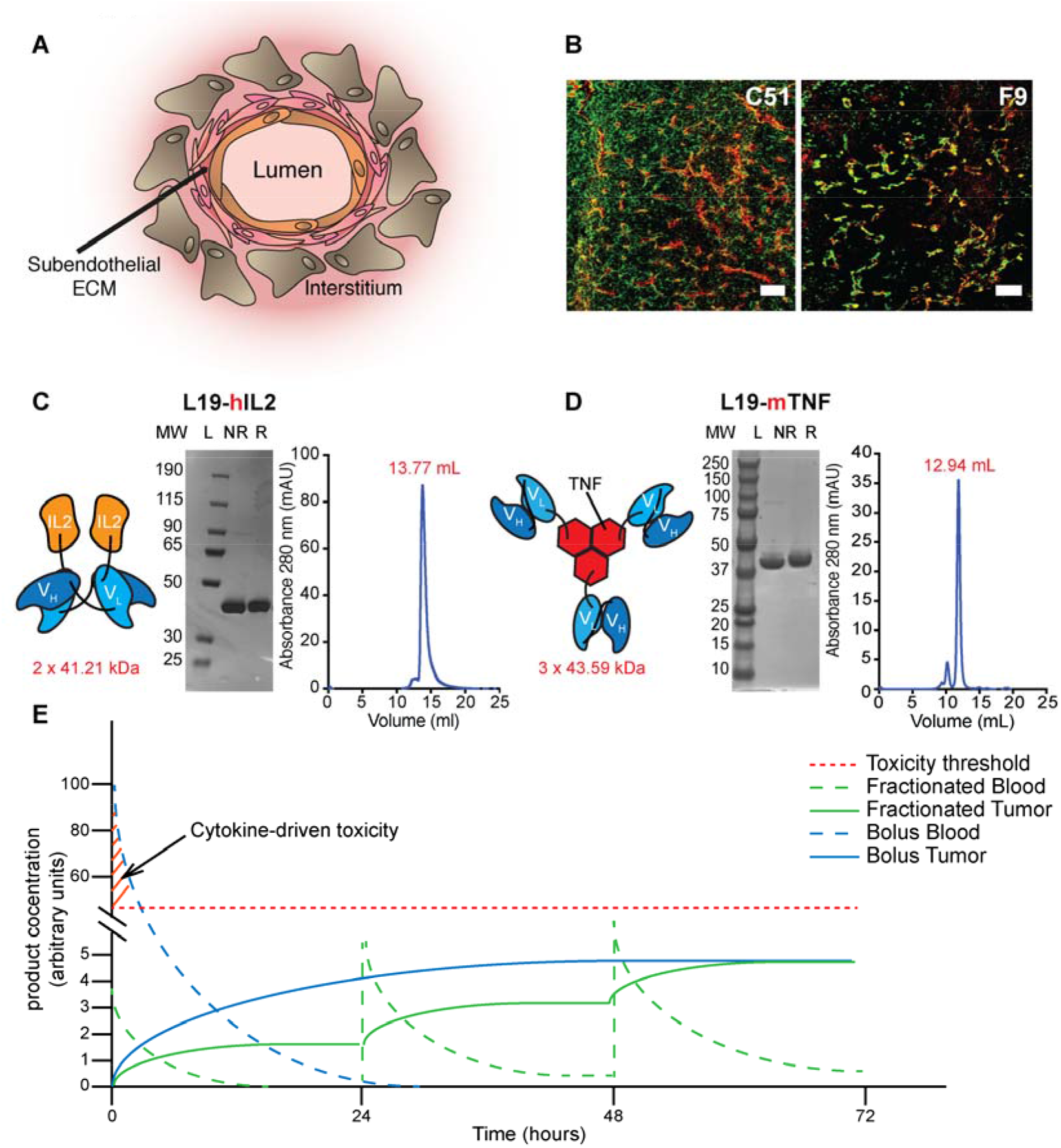
Tumor models, reagents and biodistribution rationale for using fractionated administration of antibody-cytokine fusions, rather than bolus injection. [**A**] Illustration of possible patterns of EDB in mouse models of cancer. EDB can be expressed in the basement membrane of tumor blood vessels and/or in the tumor interstitium and stroma. [**B**] Fluorescence analysis of EDB expression on C51 [**B, left panel**] and F9 [**B, right panel**] tumor sections. Cryosections were stained with L19-IgG FITC (green, AlexaFluor 488) and anti-CD31 (red, AlexaFluor 594). 10× magnification, scale bars = 100 μm. [**C, D**] Starting from left: schematic representation of the domain assembly (with mass based on amino acid sequence), SDS-PAGE analysis (MW: molecular weight, NR: non-reducing conditions, R: reducing conditions) and size exclusion chromatography profile (with retention volumes) of L19-hIL2 and L19-mTNF, respectively. [**E**] Schematic representation of the analysis reported in this study.

**Figure 1E** depicts a biodistribution rationale for using intravenous fractionated administration of antibody-cytokine fusions, rather than intravenous bolus injection. Biodistribution results may be displayed in terms of product concentration in tumor masses, blood or normal organs over time. Upon i.v. bolus injection adverse events (e.g., flu-like symptoms, hypotension) may occur as blood concentrations of the therapeutic agent stand above a critical threshold. Blood levels of drugs then typically decrease in a mono- or bi-exponential manner [58], alongside with the progressive disappearance of side effects. [30,31]. In parallel, the concentration of product in the solid tumor mass increases over time, in a process that slows down with the progressive disappearance of antibody-cytokine fusion from circulation. Products targeting stable antigens (e.g., components of the modified tumor extracellular matrix) exhibit long residence times within the neoplastic mass [59,60]. It is therefore conceivable that *by fractionating the* administration of antibody-cytokine fusions, we may achieve a *progressive* build-up of product concentration within the solid tumor mass, while keeping blood levels below a critical threshold.

We tested fractionated administration strategies in quantitative biodistribution studies with radio-labeled preparations of L19-hIL2 and of L19-mTNF in immunocompetent mice, bearing murine F9 teratocarcinomas [**Figure 2**] or C51 adenocarcinomas [**Supplementary Figure 2**], according to the experimental scheme of **Figure 2A** **and** **C**. The percent injected dose per gram of tissue or body fluid (%ID/g) for L19-hIL2, obtained by bolus injection 24 hour prior to sacrifice or by fractionated administration over time, are displayed in **Figure 2B**. As expected, comparable doses in the tumor were observed for single or fractionated (2- or 3-times) injection modalities. Increased tumor:organ ratios were observed when product injection was divided in three steps over a period of 72 hours [**Figure 2B**]. A different profile was observed for L19-mTNF. In that case, a progressive decrease in the cumulative %ID/g value in the tumor mass was observed [**Figure 2D**]. The targeted delivery of TNF to the tumor blood vessels may cause a rapid and selective shutdown of the vasculature, leading to hemorrhagic necrosis [25,36,37,61]. The conversion of a tumor mass into a necrotic scab within the first few hours is likely to prevent product accumulation for subsequent injections.

**Figure 2:**
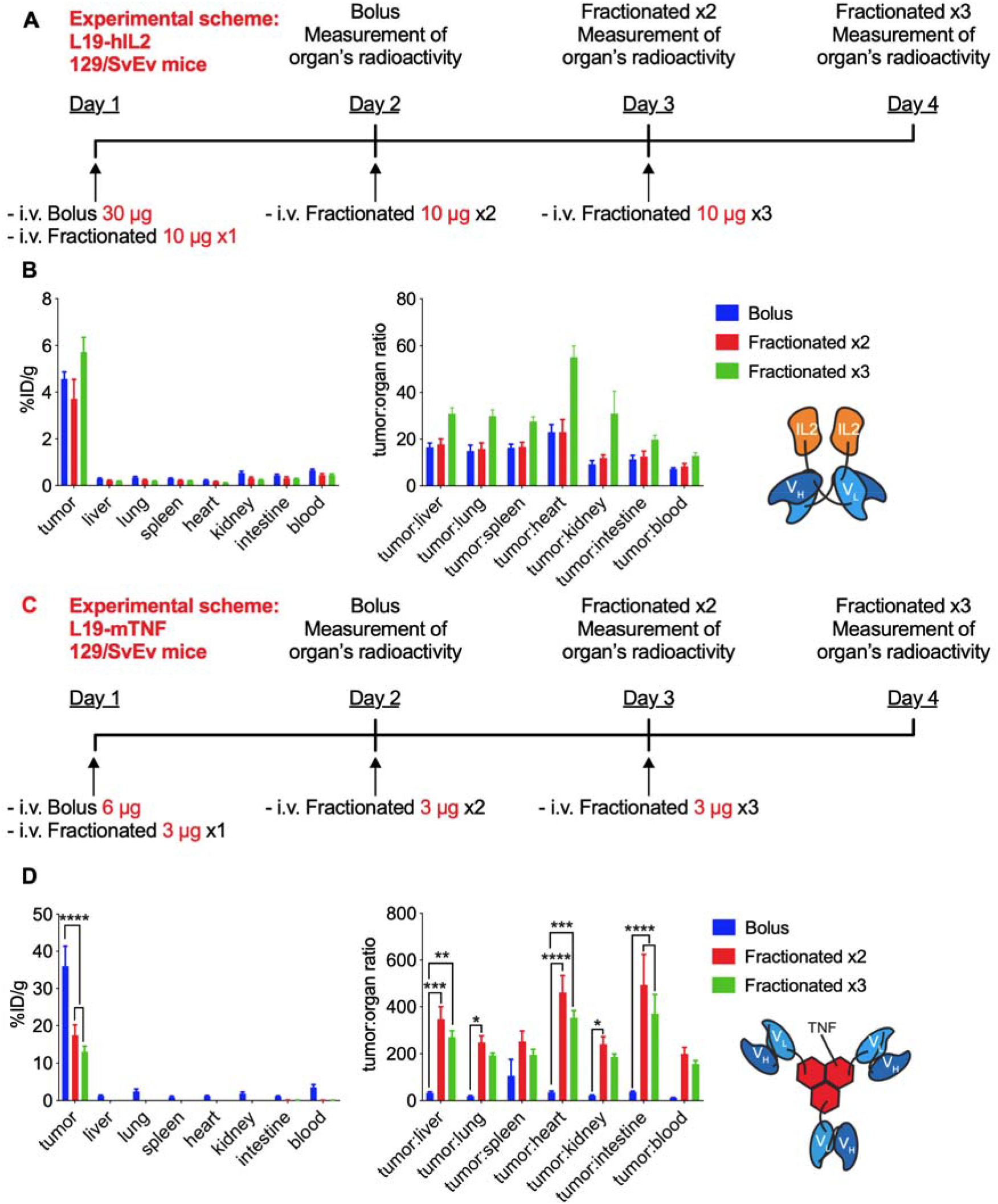
Biodistribution experiments of fractionated administration of antibody-cytokine fusions versus bolus injection. [**A**] and [**C]** Experimental schemes of the study of L19-hIL2 and L19-mTNF, respectively. Comparative quantitative biodistrubution analysis of radio-iodinated L19-hIL2 [**B**] and L19-mTNF [**D**] in immunocompetent mice bearing F9 teratocarcinoma tumors. For the bolus treatment group mice were sacrificed 24 hours after the bolus injection. For the fractionated treatment group mice were injected every 24 hours and sacrificed 24 hours after the last injection. Statistical differences were assessed between mice receiving bolus and fractionated injections of L19-mTNF or L19-hIL2. *, p<0.05; **, p<0.01 ***, p<0.001; ****, p<0.0001 (regular two-way ANOVA test with Bonferroni post-test). Results are expressed as percentage of injected dose per gram of tissue (%ID/g ± SEM), (n = 5 mice per group). For L19-hIL2 data were corrected for tumor growth correction.

We tested the antitumor-activity for L19-hIL2 in immunocompetent mice, bearing F9 teratocarcinomas [**Figure 3A**]. Unlike the human situation, IL2 shows minimal toxicity in mice. No body weight loss could be observed upon bolus administration of 100 μg and 400 μg L19-hIL2. The higher dose was efficacious in controlling tumor size for a long time period, but the lack of toxicity prevented a comparative evaluation of bolus vs. fractionated administration strategies. We repeated therapy experiments, comparing a single bolus injection of 400 μg L19-hIL2 with a fractionated administration of four doses of 200 μg, leading to a total cumulative dose of 800 μg. The latter regimen yielded a better control of tumor growth, even though complete regressions were not observed [**Figure 3B**].

**Figure 3:**
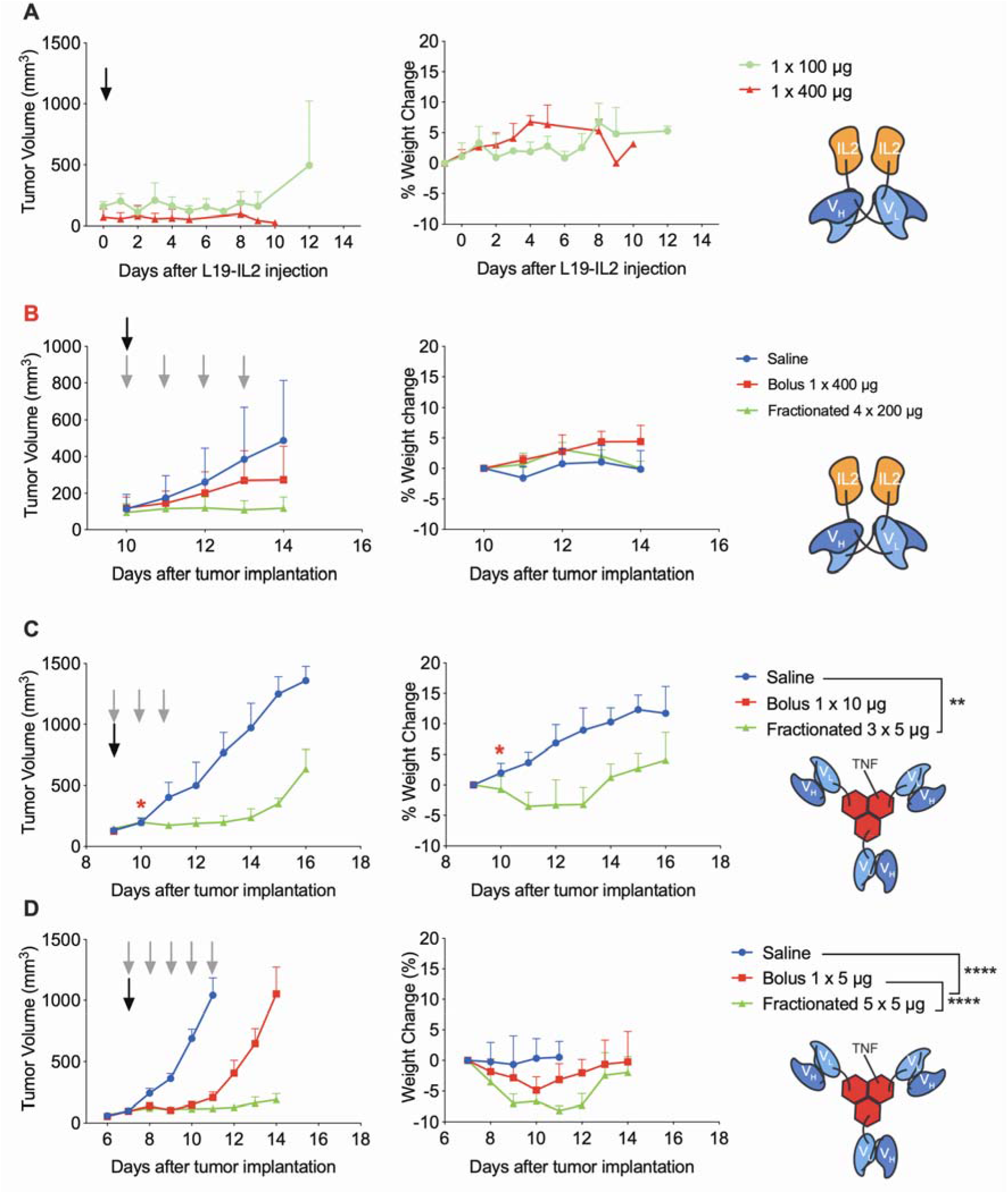
Tumor therapy experiments in 129/SvEv mice bearing F9 teratocarcinomas. [**A**] Therapy with L19-hIL2, the fusion protein was administered at 100 μg and 400 μg once (black arrow). [**B**] Comparison between a single bolus injection of 400 μg L19-hIL2 (black arrow) and four injections of 200 μg of the same product (grey arrows). While body weight profiles were excellent for all treatment groups, mice in the 4×200 μg group exhibited signs of discomfort after the last injection. [**C**] First therapy with L19-mTNF, the fusion protein was administered either as a single 10 μg bolus injection (black arrow) or as a 5 μg fractionated dose (grey arrows), which was repeated three times every 24 hours. * 5/5 mice died after the injection of 10 μg of L19-mTNF. [**D**] Second therapy with L19-mTNF, the fusion protein was administered either as a single 5 μg bolus injection (black arrow) or as a 5 μg fractionated dose (grey arrows), which was repeated five times every 24 hours. Statistical differences were assessed between mice receiving bolus and fractionated injections of L19-mTNF. **, p<0.01; ****, p<0.0001 (regular two-way ANOVA test with Bonferroni post-test). Data represent mean tumor volume ± SEM (left) and % mean body weight change ± SEM (right).

We then tested L19-mTNF in F9 teratocarcinomas in 129/SvEv mice and in C51 colorectal tumors in BALB/c mice. Different immunocompetent mouse strains exhibit different sensitivity to cytokine therapeutics. Dose fractionation allowed the safe administration of a cumulative dose of 15 μg in the F9 model, while 10 μg of L19-mTNF were lethal when administered as a bolus injection [**Figure 3C**]. Motivated by these encouraging results, we further extended dose fractionation to five consecutive injections, leading to the administration of a cumulative dose of 25 μg of product and to the prolonged inhibition of tumor growth [**Figure 3D**]. Analysis of cytokine concentrations in plasma for the different treatment groups did not explain differences in toxicity, as in 129/SvEv mice similar plasma concentrations of IL4, IL2, IL10, IL4, IL17. MIG cytokine and TGF-ß1 were observed after bolus injection (5 μg) or fractionated administration (3×3 μg) of L19-mTNF [**Figure 4**]. Pathological analysis of tumor-bearing mice after therapy experiments did not reveal macroscopic or microscopic abnormalities [**Supplementary Figure 4**].

**Figure 4:**
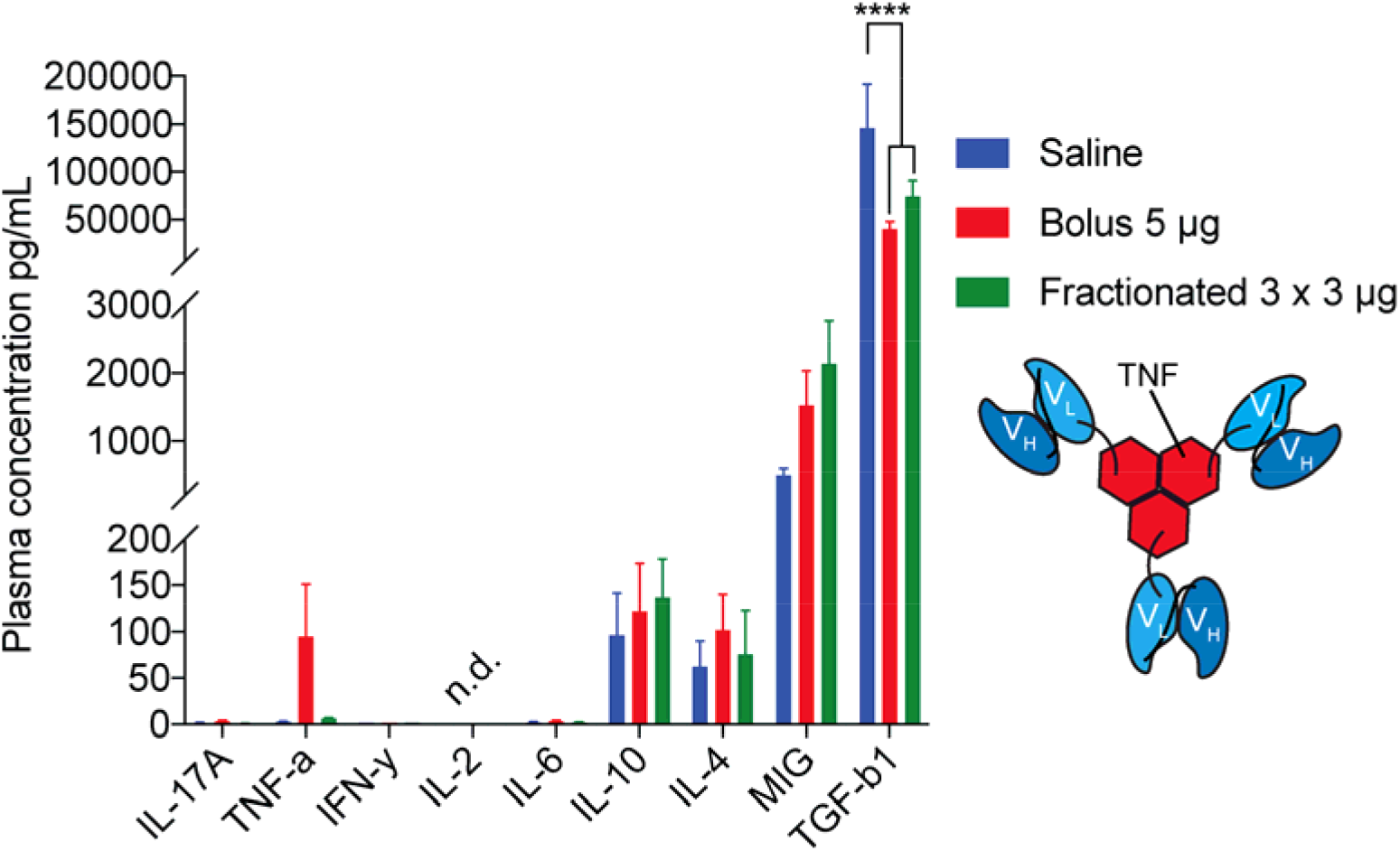
Plasma concentration of cytokines in F9 tumor bearing mice injected with either PBS, 5 μg L19-mTNF once or 3 μg L19-mTNF three times every 24 hours. Blood samples were collected 24h after the last injection. Statistical differences were assessed between mice receiving bolus and fractionated injections of L19-mTNF. ****, p<0.0001 (regular two-way ANOVA test with Bonferroni post-test). Results are expressed as plasma concentrations (pg/mL ± SEM), (n = 5 mice per group).

Dose fractionation of L19-mTNF in C51 tumors led to an opposite outcome compared to what had been observed in F9 teratocarcinomas. The product was administered either as a single 3 μg bolus injection or as 1 μg or 1.5 μg fractionated doses [**Figure 5**]. In this setting, a bolus administration of L19-mTNF showed a better anti-cancer activity compared to saline and to both fractionated treatment groups.

**Figure 5:**
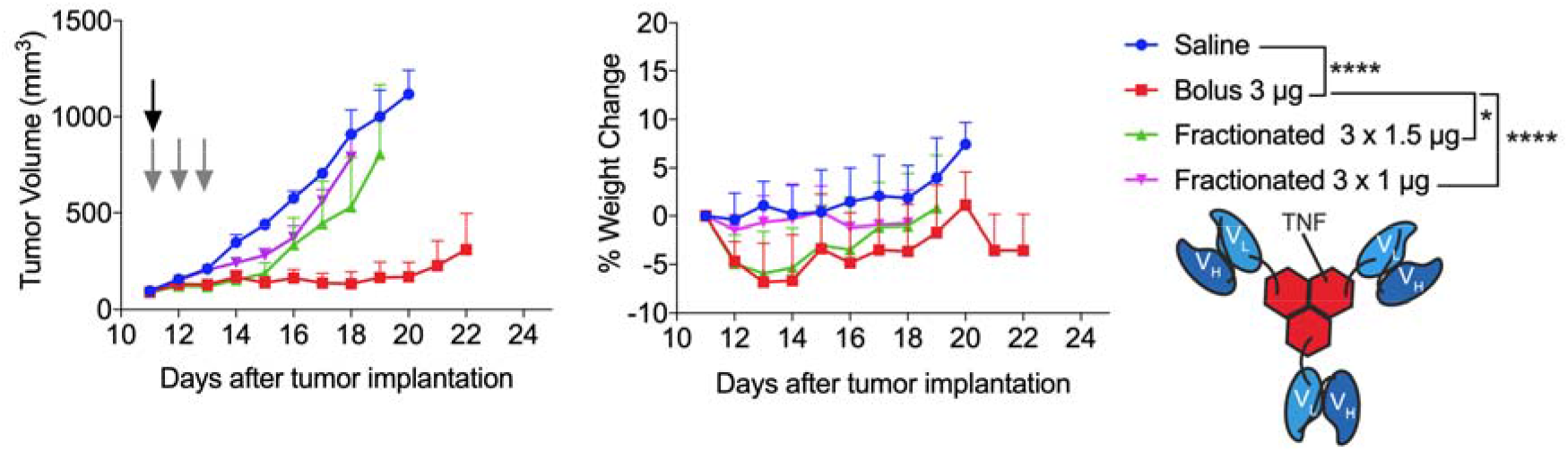
Tumor therapy experiments in BALB/c mice bearing C51 lesions. L19-mTNF was administered either as a single 3 μg bolus injection (black arrow) or as a 1 μg or 1.5 μg fractionated dose (grey arrows), which was repeated three times every 24 hours. Statistical differences were assessed between mice receiving bolus and fractionated injections of L19-mTNF. *, p<0.05; ****, p<0.0001 (regular two-way ANOVA test with Bonferroni post-test). Data represent mean tumor volume ± SEM (left) and % mean body weight change ± SEM (right).

Metronomic administration procedures have been extensively studied for many types of chemotherapeutic drugs, but are less characterized for cytokine-based biopharmaceuticals [62–64]. The therapeutic activity of NKTR-214 (a PEGylated derivative of IL2) in mice [22] and in patients [65] has been attributed to its ability to release free IL2 in a controlled fashion. The product is administered intravenously, the PEG chains are slowly hydrolyzed and as a consequence the drug becomes active. This strategy reduces systemic toxicities by providing minimal exposure of the free IL2, while achieving, over a period of 5 days, a 380-fold increased tumor exposure of the active cytokine, compared to the unmodified IL2 (i.e., adesleukin).

Certain immunotherapeutics work better when given by continuous infusion. Blinatumomab (a bispecific antibody product in BiTE™ format) has received marketing authorization for the treatment of adult and pediatric patients with B-cell precursor acute lymphoblastic leukemia (ALL). The product is administered as 30-day continuous infusion thanks to the use of an infusion pump [66]. The same agent was found to be substantially inefficacious when administered as bolus injection [67,68].

Subcutaneous injections may lead to a sustained release of immunotherapeutics. Recombinant human IL2 has shown comparable activity (but different tolerability) when given as subcutaneous administration or as bolus injection [69]. An engineered version of human interleukin-15 ALT-803) has recently shown superior biodistribution and activity profiles when administered as subcutaneous injections, while a rapid disappearance from circulation and elevated toxicity was reported for bolus intravenous administration [70].

From a theoretical viewpoint, we would expect dose fractionation to be associated with a better tolerability profile and, indeed, this feature was observed for L19-mTNF in murine models of cancer. However, the different therapeutic performance of this product in F9 and C51 tumors, upon bolus and fractionated administration [**Figures 3 and 5**], suggests that dose and schedule of L19-mTNF may ultimately require an optimization for individual indications at the clinical level.

## Acknowledgments

We would like to thank Dr. Teresa Hemmerle and Dr. Marco Colpo for helpful discussions.

## References

[1] P. Sharma, J.P. Allison, Immune checkpoint targeting in cancer therapy: Toward combination strategies with curative potential, Cell. (2015). doi:10.1016/j.cell.2015.03.030.

[2] S.C. Wei, J.H. Levine, A.P. Cogdill, Y. Zhao, N.A.A.S. Anang, M.C. Andrews, P. Sharma, J. Wang, J.A. Wargo, D. Pe’er, J.P. Allison, Distinct Cellular Mechanisms Underlie Anti-CTLA-4 and Anti-PD-1 Checkpoint Blockade, Cell. (2017). doi:10.1016/j.cell.2017.07.024.

[3] C. Robert, L. Thomas, I. Bondarenko, S. O’Day, J. Weber, C. Garbe, C. Lebbe, J.-F. Baurain, A. Testori, J.-J. Grob, N. Davidson, J. Richards, M. Maio, A. Hauschild, W.H. Miller, P. Gascon, M. Lotem, K. Harmankaya, R. Ibrahim, S. Francis, T.-T. Chen, R. Humphrey, A. Hoos, J.D. Wolchok, Ipilimumab plus dacarbazine for previously untreated metastatic melanoma., N. Engl. J. Med. (2011). doi:10.1056/NEJMoa1104621.

[4] C. Robert, G. V Long, B. Brady, C. Dutriaux, M. Maio, L. Mortier, J.C. Hassel, P. Rutkowski, C. McNeil, E. Kalinka-Warzocha, K.J. Savage, M.M. Hernberg, C. Lebbé, J. Charles, C. Mihalcioiu, V. Chiarion-Sileni, C. Mauch, F. Cognetti, A. Arance, H. Schmidt, D. Schadendorf, H. Gogas, L. Lundgren-Eriksson, C. Horak, B. Sharkey, I.M. Waxman, V. Atkinson, P.A. Ascierto, Nivolumab in previously untreated melanoma without BRAF mutation., N. Engl. J. Med. (2015). doi:10.1056/NEJMoa1412082.

[5] C. Robert, J. Schachter, G. V Long, A. Arance, J.J. Grob, L. Mortier, A. Daud, M.S. Carlino, C. McNeil, M. Lotem, J. Larkin, P. Lorigan, B. Neyns, C.U. Blank, O. Hamid, C. Mateus, R. Shapira-Frommer, M. Kosh, H. Zhou, N. Ibrahim, S. Ebbinghaus, A. Ribas, KEYNOTE-006 investigators, Pembrolizumab versus Ipilimumab in Advanced Melanoma., N. Engl. J. Med. (2015). doi:10.1056/NEJMoa1503093.

[6] J. Larkin, V. Chiarion-Sileni, R. Gonzalez, J.J. Grob, C.L. Cowey, C.D. Lao, D. Schadendorf, R. Dummer, M. Smylie, P. Rutkowski, P.F. Ferrucci, A. Hill, J. Wagstaff, M.S. Carlino, J.B. Haanen, M. Maio, I. Marquez-Rodas, G.A. McArthur, P.A. Ascierto, G. V Long, M.K. Callahan, M.A. Postow, K. Grossmann, M. Sznol, B. Dreno, L. Bastholt, A. Yang, L.M. Rollin, C. Horak, F.S. Hodi, J.D. Wolchok, Combined Nivolumab and Ipilimumab or Monotherapy in Untreated Melanoma., N. Engl. J. Med. (2015). doi:10.1056/NEJMoa1504030.

[7] H.L. Kaufman, J. Russell, O. Hamid, S. Bhatia, P. Terheyden, S.P. D’Angelo, K.C. Shih, C. Lebbé, G.P. Linette, M. Milella, I. Brownell, K.D. Lewis, J.H. Lorch, K. Chin, L. Mahnke, A. von Heydebreck, J.M. Cuillerot, P. Nghiem, Avelumab in patients with chemotherapy-refractory metastatic Merkel cell carcinoma: a multicentre, single-group, open-label, phase 2 trial, Lancet Oncol. (2016). doi:10.1016/S1470-2045(16)30364-3.

[8] L. Fehrenbacher, A. Spira, M. Ballinger, M. Kowanetz, J. Vansteenkiste, J. Mazieres, K. Park, D. Smith, A. Artal-Cortes, C. Lewanski, F. Braiteh, D. Waterkamp, P. He, W. Zou, D.S. Chen, J. Yi, A. Sandler, A. Rittmeyer, Atezolizumab versus docetaxel for patients with previously treated non-small-cell lung cancer (POPLAR): A multicentre, open-label, phase 2 randomised controlled trial, Lancet. (2016). doi:10.1016/S0140-6736(16)00587-0.

[9] D. Planchard, T. Yokoi, M.J. McCleod, J.R. Fischer, Y.C. Kim, M. Ballas, K. Shi, J.C. Soria, A Phase III Study of Durvalumab (MEDI4736) with or Without Tremelimumab for Previously Treated Patients with Advanced NSCLC: Rationale and Protocol Design of the ARCTIC Study, Clin. Lung Cancer. (2016). doi:10.1016/j.cllc.2016.03.003.

[10] F. Bootz, D. Neri, Immunocytokines: A novel class of products for the treatment of chronic inflammation and autoimmune conditions, Drug Discov. Today. (2016). doi:10.1016/j.drudis.2015.10.012.

[11] N. Pasche, D. Neri, Immunocytokines: A novel class of potent armed antibodies, Drug Discov. Today. (2012). doi:10.1016/j.drudis.2012.01.007.

[12] S.A. Rosenberg, IL-2: The First Effective Immunotherapy for Human Cancer, J. Immunol. (2014). doi:10.4049/jimmunol.1490019.

[13] J.A. Klapper, S.G. Downey, F.O. Smith, J.C. Yang, M.S. Hughes, U.S. Kammula, R.M. Sherry, R.E. Royal, S.M. Steinberg, S. Rosenberg, High-dose interleukin-2 for the treatment of metastatic renal cell carcinoma: A retrospective analysis of response and survival in patients treated in the Surgery Branch at the National Cancer Institute between 1986 and 2006, Cancer. (2008). doi:10.1002/cncr.23552.

[14] F.O. Smith, S.G. Downey, J.A. Klapper, J. C. Yang, R.M. Sherry, R.E. Royal, U.S. Kammula, M.S. Hughes, N.P. Restifo, C.L. Levy, D. E. White, S.M. Steinberg, S.A. Rosenberg, Treatment of metastatic melanoma using Interleukin-2 alone or in conjunction with vaccines, Clin. Cancer Res. (2008). doi:10.1158/1078-0432.CCR-08-0116.

[15] S.A. Rosenberg, M.T. Lotze, L.M. Muul, A.E. Chang, F.P. Avis, S. Leitman, W.M. Linehan, C.N. Robertson, R.E. Lee, J.T. Rubin, C.A. Seipp, C.G. Simpson, D.E. White, A Progress Report on the Treatment of 157 Patients with Advanced Cancer Using Lymphokine-Activated Killer Cells and Interleukin-2 or High-Dose Interleukin-2 Alone, N. Engl. J. Med. (2010). doi:10.1056/nejm198704093161501.

[16] R. van Horssen, TNF- in Cancer Treatment: Molecular Insights, Antitumor Effects, and Clinical Utility, Oncologist. (2006). doi:10.1634/theoncologist.11-4-397.

[17] C. Verhoef, J.H.W. Wilt, D.J. Grünhagen, A.N. Geel, T.L.M. Hagen, A.M.M. Eggermont, Isolated limb perfusion with melphalan and TNF-α in the treatment of extremity sarcoma, Curr. Treat. Options Oncol. (2007). doi:10.1007/s11864-007-0044-y.

[18] A.M.M. Eggermont, H.S. Koops, J.M. Klausner, B.B.R. Kroon, P.M. Schlag, D. Liénard, A.N. Van Geel, H.J. Hoekstra, I. Meller, O.E. Nieweg, C. Kettelhack, G. Ben-Ari, J.C. Pector, F.J. Lejeune, Isolated limb perfusion with tumor necrosis factor and melphalan for limb salvage in 186 patients with locally advanced soft tissue extremity sarcomas: The cumulative multicenter European experience, Ann. Surg. (1996). doi:10.1097/00000658-199612000-00011.

[19] J. Jakob, P. Hohenberger, Role of isolated limb perfusion with recombinant human tumor necrosis factor α and melphalan in locally advanced extremity soft tissue sarcoma, Cancer. (2016). doi:10.1002/cncr.29991.

[20] H. Strander, S. Einhorn, Interferon therapy in neoplastic diseases., Philos. Trans. R. Soc. Lond. B. Biol. Sci. (1982). doi:10.1098/rstb.1982.0111.

[21] H. Mellstedt, M. Björkholm, B. Johansson, A. Ahre, G. Holm, H. Strander, Interferon Therapy In Myelomatosis, Lancet. (1979). doi:10.1016/S0140-6736(79)90770-0.

[22] D.H. Charych, U. Hoch, J.L. Langowski, S.R. Lee, M.K. Addepalli, P.B. Kirk, D. Sheng, X. Liu, P.W. Sims, L.A. VanderVeen, C.F. Ali, T.K. Chang, M. Konakova, R.L. Pena, R.S. Kanhere, Y.M. Kirksey, C. Ji, Y. Wang, J. Huang, T.D. Sweeney, S.S. Kantak, S.K. Doberstein, NKTR-214, an Engineered Cytokine with Biased IL2 Receptor Binding, Increased Tumor Exposure, and Marked Efficacy in Mouse Tumor Models, Clin. Cancer Res. (2016). doi:10.1158/1078-0432.CCR-15-1631.

[23] D. Neri, P.M. Sondel, Immunocytokines for cancer treatment: Past, present and future, Curr. Opin. Immunol. (2016). doi:10.1016/j.coi.2016.03.006.

[24] D. Neri, Antibody–Cytokine Fusions: Versatile Products for the Modulation of Anticancer Immunity, Cancer Immunol. Res. (2019). doi:10.1158/2326-6066.CIR-18-0622.

[25] L. Borsi, E. Balza, B. Carnemolla, F. Sassi, P. Castellani, A. Berndt, H. Kosmehl, A. Birò, A. Siri, P. Orecchia, J. Grassi, D. Neri, L. Zardi, Selective targeted delivery of TNFα to tumor blood vessels, Blood. (2003). doi:10.1182/blood-2003-04-1039.

[26] B. Carnemolla, L. Borsi, E. Balza, P. Castellani, R. Meazza, A. Berndt, S. Ferrini, H. Kosmehl, D. Neri, L. Zardi, Enhancement of the antitumor properties of interleukin-2 by its targeted delivery to the tumor blood vessel extracellular matrix, Blood. (2002). doi:10.1182/blood.V99.5.1659.

[27] C. Halin, S. Rondini, F. Nilsson, A. Berndt, H. Kosmehl, L. Zardi, D. Neri, Enhancement of the antitumor activity of interleukin-12 by targeted delivery to neovasculature, Nat. Biotechnol. (2002). doi:10.1038/nbt0302-264.

[28] R. Danielli, R. Patuzzo, A.M. Di Giacomo, G. Gallino, A. Maurichi, A. Di Florio, O. Cutaia, A. Lazzeri, C. Fazio, C. Miracco, L. Giovannoni, G. Elia, D. Neri, M. Maio, M. Santinami, Intralesional administration of L19-IL2/L19-TNF in stage III or stage IVM1a melanoma patients: Results of a phase II study, Cancer Immunol. Immunother. (2015). doi:10.1007/s00262-015-1704-6.

[29] B. Weide, T.K. Eigentler, A. Pflugfelder, H. Zelba, A. Martens, G. Pawelec, L. Giovannoni, P.A. Ruffini, G. Elia, D. Neri, R. Gutzmer, J.C. Becker, C. Garbe, Intralesional Treatment of Stage III Metastatic Melanoma Patients with L19-IL2 Results in Sustained Clinical and Systemic Immunologic Responses, Cancer Immunol. Res. (2014). doi:10.1158/2326-6066.cir-13-0206.

[30] T.K. Eigentler, B. Weide, F. De Braud, G. Spitaleri, A. Romanini, A. Pflugfelder, R. Gonzaĺez-Iglesias, A. Tasciotti, L. Giovannoni, K. Schwager, V. Lovato, M. Kaspar, E. Trachsel, H.D. Menssen, D. Neri, C. Garbe, A dose-escalation and signal-generating study of the immunocytokine L19-IL2 in combination with dacarbazine for the therapy of patients with metastatic melanoma, Clin. Cancer Res. (2011). doi:10.1158/1078-0432.CCR-11-1203.

[31] M. Johannsen, G. Spitaleri, G. Curigliano, J. Roigas, S. Weikert, C. Kempkensteffen, A. Roemer, C. Kloeters, P. Rogalla, G. Pecher, K. Miller, A. Berndt, H. Kosmehl, E. Trachsel, M. Kaspar, V. Lovato, R. González-Iglesias, L. Giovannoni, H.D. Menssen, D. Neri, F. De Braud, The tumour-targeting human L19-IL2 immunocytokine: Preclinical safety studies, phase i clinical trial in patients with solid tumours and expansion into patients with advanced renal cell carcinoma, Eur. J. Cancer. (2010). doi:10.1016/j.ejca.2010.07.033.

[32] G. Spitaleri, R. Berardi, C. Pierantoni, T. De Pas, C. Noberasco, C. Libbra, R. González-Iglesias, L. Giovannoni, A. Tasciotti, D. Neri, H.D. Menssen, F. De Braud, Phase I/II study of the tumour-targeting human monoclonal antibody-cytokine fusion protein L19-TNF in patients with advanced solid tumours, J. Cancer Res. Clin. Oncol. (2013). doi:10.1007/s00432-012-1327-7.

[33] F. Papadia, V. Basso, R. Patuzzo, A. Maurichi, A. Di Florio, L. Zardi, E. Ventura, R. González-Iglesias, V. Lovato, L. Giovannoni, A. Tasciotti, D. Neri, M. Santinami, H.D. Menssen, F. De Cian, Isolated limb perfusion with the tumor-targeting human monoclonal antibody-cytokine fusion protein L19-TNF plus melphalan and mild hyperthermia in patients with locally advanced extremity melanoma, J. Surg. Oncol. (2013). doi:10.1002/jso.23168.

[34] F. Viti, L. Tarli, L. Giovannoni, L. Zardi, D. Neri, Increased binding affinity and valence of recombinant antibody fragments lead to improved targeting of tumoral angiogenesis, Cancer Res. (1999).

[35] T. Hemmerle, P. Probst, L. Giovannoni, A.J. Green, T. Meyer, D. Neri, The antibody-based targeted delivery of TNF in combination with doxorubicin eradicates sarcomas in mice and confers protective immunity, Br. J. Cancer. (2013). doi:10.1038/bjc.2013.421.

[36] R. De Luca, A. Soltermann, F. Pretto, C. Pemberton-Ross, G. Pellegrini, S. Wulhfard, D. Neri, Potency-matched Dual Cytokine–Antibody Fusion Proteins for Cancer Therapy, Mol. Cancer Ther. (2017). doi:10.1158/1535-7163.mct-17-0211.

[37] P. Probst, J. Kopp, A. Oxenius, M.P. Colombo, D. Ritz, T. Fugmann, D. Neri, Sarcoma eradication by doxorubicin and targeted TNF relies upon CD8 + T-cell recognition of a retroviral antigen, Cancer Res. (2017). doi:10.1158/0008-5472.CAN-16-2946.

[38] C. Hutmacher, N. Gonzalo Núñez, A.R. Liuzzi, B. Becher, D. Neri, Targeted Delivery of IL2 to the Tumor Stroma Potentiates the Action of Immune Checkpoint Inhibitors by Preferential Activation of NK and CD8 + T Cells, Cancer Immunol. Res. (2019). doi:10.1158/2326-6066.cir-18-0566.

[39] F. Bensch, M.M. Smeenk, S.C. van Es, J.R. de Jong, C.P. Schröder, S.F. Oosting, M.N. Lub-de Hooge, C.W. Menke-van der Houven van Oordt, A.H. Brouwers, R. Boellaard, E.G.E. de Vries, Comparative biodistribution analysis across four different 89Zr-monoclonal antibody tracers-The first step towards an imaging warehouse, Theranostics. (2018). doi:10.7150/thno.26370.

[40] M.R. Buist, P. Kenemans, C.F.M. Molthoff, J.C. Roos, W. Den Hollander, M. Brinkhuis, J.P.A. Baak, Tumor uptake of intravenously administered radiolabeled antibodies in ovarian carcinoma patients in relation to antigen expression and other tumor characteristics, Int. J. Cancer. (1995). doi:10.1002/ijc.2910640204.

[41] S.D. Gillies, A new platform for constructing antibody-cytokine fusion proteins (immunocytokines) with improved biological properties and adaptable cytokine activity, Protein Eng. Des. Sel. (2013). doi:10.1093/protein/gzt045.

[42] D. Venetz, D. Koovely, B. Weder, D. Neri, Targeted reconstitution of cytokine activity upon antigen binding using split cytokine antibody fusion proteins, J. Biol. Chem. (2016). doi:10.1074/jbc.M116.737734.

[43] S.D. Gillies, Y. Lan, B. Brunkhorst, W.K. Wong, Y. Li, K.M. Lo, Bi-functional cytokine fusion proteins for gene therapy and antibody-targeted treatment of cancer, Cancer Immunol. Immunother. (2002). doi:10.1007/s00262-002-0302-6.

[44] R. De Luca, D. Neri, Potentiation of PD-L1 blockade with a potency-matched dual cytokine–antibody fusion protein leads to cancer eradication in BALB/c-derived tumors but not in other mouse strains, Cancer Immunol. Immunother. (2018). doi:10.1007/s00262-018-2194-0.

[45] R. De Luca, P. Kachel, K. Kropivsek, B. Snijder, M.G. Manz, D. Neri, A novel dual-cytokine-antibody fusion protein for the treatment of CD38-positive malignancies, Protein Eng. Des. Sel. (2018). doi:10.1093/protein/gzy015.

[46] C. Halin, V. Gafner, M.E. Villani, L. Borsi, A. Berndt, H. Kosmehl, L. Zardi, D. Neri, Synergistic therapeutic effects of a tumor targeting antibody fragment, fused to interleukin 12 and to Tumor Necrosis Factor α, Cancer Res. (2003).

[47] V. Kermer, N. Hornig, M. Harder, A. Bondarieva, R.E. Kontermann, D. Muller, Combining Antibody-Directed Presentation of IL-15 and 4-1BBL in a Trifunctional Fusion Protein for Cancer Immunotherapy, Mol. Cancer Ther. (2014). doi:10.1158/1535-7163.mct-13-0282.

[48] N. Beha, M. Harder, S. Ring, R.E. Kontermann, D. Müller, IL-15-based trifunctional antibody-fusion proteins with costimulatory TNF-superfamily ligands in the single-chain format for cancer immunotherapy, Mol. Cancer Ther. (2019). doi:10.1158/1535-7163.mct-18-1204.

[49] C. Klein, I. Waldhauer, V.G. Nicolini, A. Freimoser-Grundschober, T. Nayak, D.J. Vugts, C. Dunn, M. Bolijn, J. Benz, M. Stihle, S. Lang, M. Roemmele, T. Hofer, E. van Puijenbroek, D. Wittig, S. Moser, O. Ast, P. Brünker, I.H. Gorr, S. Neumann, M.C. de Vera Mudry, H. Hinton, F. Crameri, J. Saro, S. Evers, C. Gerdes, M. Bacac, G. van Dongen, E. Moessner, P. Umaña, Cergutuzumab amunaleukin (CEA-IL2v), a CEA-targeted IL-2 variant-based immunocytokine for combination cancer immunotherapy: Overcoming limitations of aldesleukin and conventional IL-2-based immunocytokines, Oncoimmunology. (2017). doi:10.1080/2162402X.2016.1277306.

[50] S.L. Pogue, T. Taura, M. Bi, Y. Yun, A. Sho, G. Mikesell, C. Behrens, M. Sokolovsky, H. Hallak, M. Rosenstock, E. Sanchez, H. Chen, J. Berenson, A. Doyle, S. Nock, D.S. Wilson, Targeting attenuated interferon-α to myeloma cells with a CD38 antibody induces potent tumor regression with reduced off-target activity, PLoS One. (2016). doi:10.1371/journal.pone.0162472.

[51] M.J. Bernett, R. Varma, C. Bonzon, R. Rashid, L. Bogaert, K. Liu, S. Schubbert, K.N. Avery, I.W. Leung, N. Rodriguez, S.Y. Chu, U.S. Muchhal, G.L. Moore, J.R. Desjarlais, Abstract 5565: Potency-reduced IL15/IL15Rα heterodimeric Fc-fusions display enhanced in vivo activity through increased exposure, Cancer Res. 78 (2018) 5565 LP – 5565. doi:10.1158/1538-7445.AM2018-5565.

[52] N. Arenas-Ramirez, C. Zou, S. Popp, D. Zingg, B. Brannetti, E. Wirth, T. Calzascia, J. Kovarik, L. Sommer, G. Zenke, J. Woytschak, C.H. Regnier, A. Katopodis, O. Boyman, Improved cancer immunotherapy by a CD25-mimobody conferring selectivity to human interleukin-2, Sci. Transl. Med. (2016). doi:10.1126/scitranslmed.aag3187.

[53] L. Tarli, E. Balza, F. Viti, L. Borsi, P. Castellani, D. Berndorff, L. Dinkelborg, D. Neri, L. Zardi, A high-affinity human antibody that targets tumoral blood vessels, Blood. (1999).

[54] J.N. Rybak, C. Roesli, M. Kaspar, A. Villa, D. Neri, The extra-domain A of fibronectin is a vascular marker of solid tumors and metastases, Cancer Res. (2007). doi:10.1158/0008-5472.CAN-07-1436.

[55] K. Frey, M. Fiechter, K. Schwager, B. Belloni, M.J. Barysch, D. Neri, R. Dummer, Different patterns of fibronectin and tenascin-C splice variants expression in primary and metastatic melanoma lesions, Exp. Dermatol. (2011). doi:10.1111/j.1600-0625.2011.01314.x.

[56] K. Schwager, A. Villa, C. Rösli, D. Neri, M. Rösli-Khabas, G. Moser, A comparative immunofluorescence analysis of three clinical-stage antibodies in head and neck cancer, Head Neck Oncol. (2011). doi:10.1186/1758-3284-3-25.

[57] C. Schliemann, A. Wiedmer, M. Pedretti, M. Szczepanowski, W. Klapper, D. Neri, Three clinical-stage tumor targeting antibodies reveal differential expression of oncofetal fibronectin and tenascin-C isoforms in human lymphoma, Leuk. Res. (2009). doi:10.1016/j.leukres.2009.06.025.

[58] L. Borsi, E. Balza, M. Bestagno, P. Castellani, B. Carnemolla, A. Biro, A. Leprini, J. Sepulveda, O. Burrone, D. Neri, L. Zardi, Selective targeting of tumoral vasculature: Comparison of different formats of an antibody (l19) to the ED-B domain of fibronectin, Int. J. Cancer. (2002). doi:10.1002/ijc.10662.

[59] K. Schwager, M. Kaspar, F. Bootz, R. Marcolongo, E. Paresce, D. Neri, E. Trachsel, Preclinical characterization of DEKAVIL (F8-IL10), a novel clinical-stage immunocytokine which inhibits the progression of collagen-induced arthritis, Arthritis Res. Ther. (2009). doi:10.1186/ar2814.

[60] T. Ongaro, M. Matasci, S. Cazzamalli, B. Gouyou, R. De Luca, D. Neri, A. Villa, A novel anti-cancer L19-interleukin-12 fusion protein with an optimized peptide linker efficiently localizes in vivo at the site of tumors, J. Biotechnol. (2019). doi:10.1016/j.jbiotec.2018.12.004.

[61] B. Ziffels, F. Pretto, D. Neri, Intratumoral administration of IL2-and TNF-based fusion proteins cures cancer without establishing protective immunity, Immunotherapy. (2018). doi:10.2217/imt-2017-0119.

[62] R.S. Kerbel, Y. Shaked, The potential clinical promise of ‘multimodality’ metronomic chemotherapy revealed by preclinical studies of metastatic disease, Cancer Lett. (2017). doi:10.1016/j.canlet.2017.02.005.

[63] G. Bocci, R.S. Kerbel, Pharmacokinetics of metronomic chemotherapy: A neglected but crucial aspect, Nat. Rev. Clin. Oncol. (2016). doi:10.1038/nrclinonc.2016.64.

[64] R.S. Kerbel, B.A. Kamen, The anti-angiogenic basis of metronomic chemotherapy, Nat. Rev. Cancer. (2004). doi:10.1038/nrc1369.

[65] S.-E. Bentebibel, M.E. Hurwitz, C. Bernatchez, C. Haymaker, C.W. Hudgens, H.M. Kluger, M.T. Tetzlaff, M.A. Tagliaferri, J. Zalevsky, U. Hoch, C. Fanton, S. Aung, P. Hwu, B.D. Curti, N.M. Tannir, M. Sznol, A. Diab, A First-in-Human Study and Biomarker Analysis of NKTR-214, a Novel IL-2-Receptor Beta/Gamma (βγ)-Biased Cytokine, in Patients With Advanced or Metastatic Solid Tumors, Cancer Discov. (2019). doi:10.1158/2159-8290.CD-18-1495.

[66] H. Kantarjian, A. Stein, N. Gökbuget, A.K. Fielding, A.C. Schuh, J.-M. Ribera, A. Wei, H. Dombret, R. Foà, R. Bassan, Ö. Arslan, M.A. Sanz, J. Bergeron, F. Demirkan, E. Lech-Maranda, A. Rambaldi, X. Thomas, H.-A. Horst, M. Brüggemann, W. Klapper, B.L. Wood, A. Fleishman, D. Nagorsen, C. Holland, Z. Zimmerman, M.S. Topp, Blinatumomab versus Chemotherapy for Advanced Acute Lymphoblastic Leukemia., N. Engl. J. Med. (2017). doi:10.1056/NEJMoa1609783.

[67] G. Friberg, Blinatumomab (Blincyto): Lessons learned from the bispecific t-cell engager (BiTE) in acute lymphocytic leukemia (ALL), Ann. Oncol. (2017). doi:10.1093/annonc/mdx150.

[68] D. Nagorsen, P. Kufer, P.A. Baeuerle, R. Bargou, Blinatumomab: A historical perspective, Pharmacol. Ther. (2012). doi:10.1016/j.pharmthera.2012.07.013.

[69] O. Eton, M.G. Rosenblum, S.S. Legha, W. Zhang, M.J. East, A. Bedikian, N. Papadopoulos, A. Buzaid, R.S. Benjamin, Phase I trial of subcutaneous recombinant human interleukin-2 in patients with metastatic melanoma, Cancer. (2002). doi:10.1002/cncr.10631.

[70] K. Margolin, C. Morishima, V. Velcheti, J.S. Miller, S.M. Lee, A.W. Silk, S.G. Holtan, A.M. Lacroix, S.P. Fling, J.C. Kaiser, J.O. Egan, M. Jones, P.R. Rhode, A.D. Rock, M.A. Cheever, H.C. Wong, M.S. Ernstoff, Phase I trial of ALT-803, a novel recombinant IL15 complex, in patients with advanced solid tumors, Clin. Cancer Res. (2018). doi:10.1158/1078-0432.CCR-18-0945.

