## supplementary Figure Legends for "Comparative evaluation of bolus and fractionated administration modalities for two antibody-cytokine fusions in immunocompetent tumor-bearing mice"

**A**

EVQLLESGGGLVQPGGSLRLSCAASGFTFSSFSMSWVRQAPGKGLEWVSSISGSSGTTYYADSVKGRFTISRDNSKNTLYLQMNSLRAEDTAVYYCAKPFPYFDYWGQGTLVTVSS*GDGSSGGSGGAS*EIVLTQSPGTLSLSPGERATLSCRASQSVSSSFLAWYQQKPGQAPRLLIYYASSRATGIPDRFSGSGSGTDFTLTISRLEPEDFAVYYCQQTGRIPPTFGQGTKVEIKSSSSGSSSSGSSSSGLRSSSQNSSDKPVAHVVANHQVEEQLEWLSQRANALLANGMDLKDNQLVVPADGLYLVYSQVLFKGQGCPDYVLLTHTVSRFAISYQEKVNLLSAVKSPCPKDTPEGAELKPWYEPIYLGGVFQLEKGDQLSAEVNLPKYLDFAESGQVYFGVIAL

**B**

EVQLLESGGGLVQPGGSLRLSCAASGFTFSSFSMSWVRQAPGKGLEWVSSISGSSGTTYYADSVKGRFTISRDNSKNTLYLQMNSLRAEDTAVYYCAKPFPYFDYWGQGTLVTVSS*GDGSSGGSGGAS*EIVLTQSPGTLSLSPGERATLSCRASQSVSSSFLAWYQQKPGQAPRLLIYYASSRATGIPDRFSGSGSGTDFTLTISRLEPEDFAVYYCQQTGRIPPTFGQGTKVEIKEFSSSSGSSSSGSSSSGAPTSSSTKKTQLQLEHLLLDLQMILNGINNYKNPKLTRMLTFKFYMPKKATELKHLQCLEEELKPLEEVLNLAQSKNFHLRPRDLISNINVIVLELKGSETTFMCEYADETATIVEFLNRWITFCQSIISTLT

[**Supplementary Figure 1**] Aminoacid sequences of [**A**] L19-mTNF and [**B**] L19-hIL2.

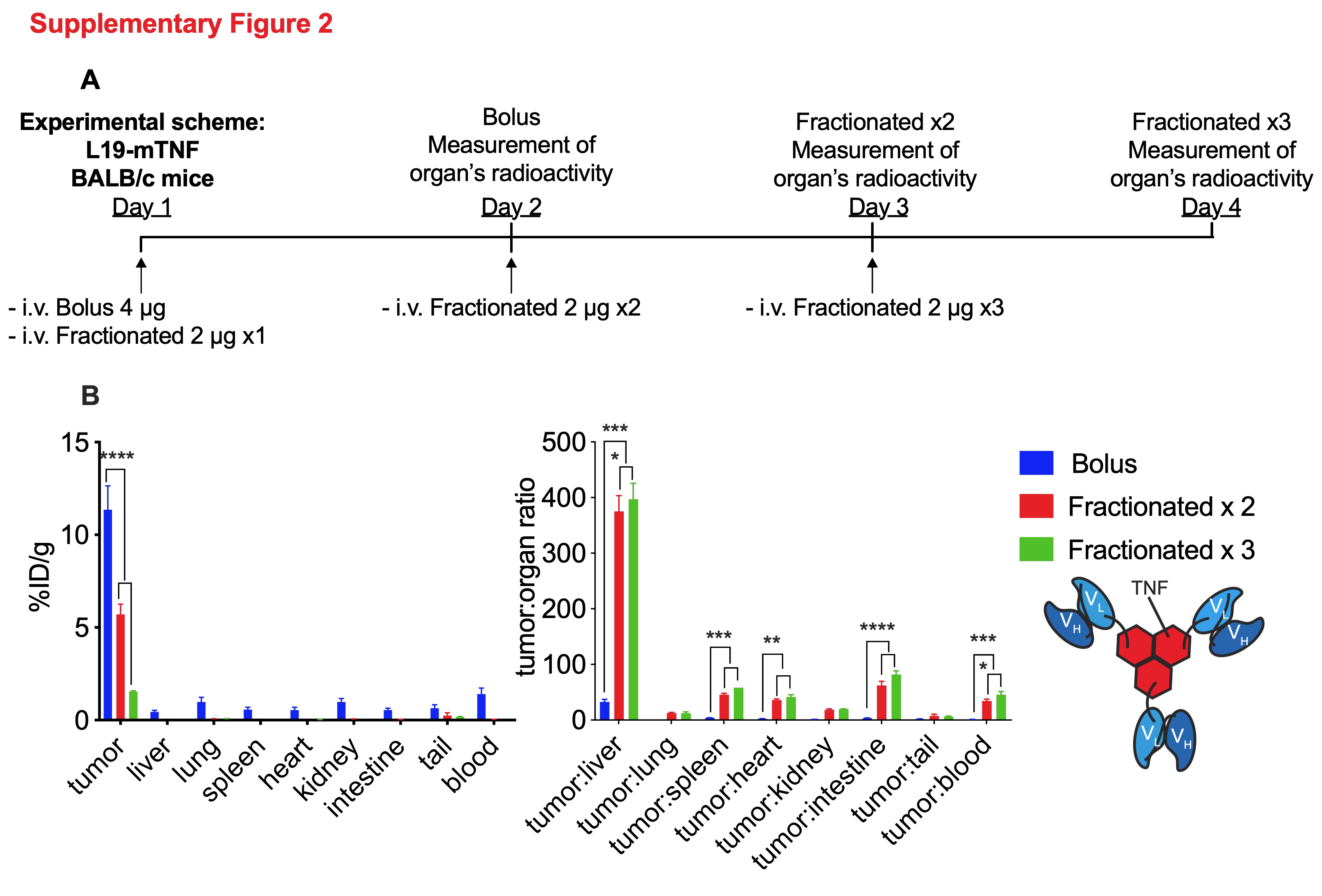

[**Supplementary Figure 2**] Biodistribution experiments of fractionated administration of L19-mTNF versus bolus injection. [**A**] Experimental schemes of the study of L19-hIL2 and L19-mTNF, respectively. Comparative quantitative biodistrubution analysis of radio-iodinated L19-mTNF [**B**] in immunocompetent mice bearing C51 Colon carcinomas. For the bolus treatment group mice were sacrificed 24 hours after the bolus injection. For the fractionated treatment group mice were injected every 24 hours and sacrificed 24 hours after the last injection. Statistical differences were assessed between mice receiving bolus and fractionated injections of L19-mTNF or L19-hIL2. *, p<0.05; **, p<0.01 ***, p<0.001; ****, p<0.0001 (regular two-way ANOVA test with Bonferroni post-test). Results are expressed as percentage of injected dose per gram of tissue (%ID/g ± SEM), (n = 5 mice per group).

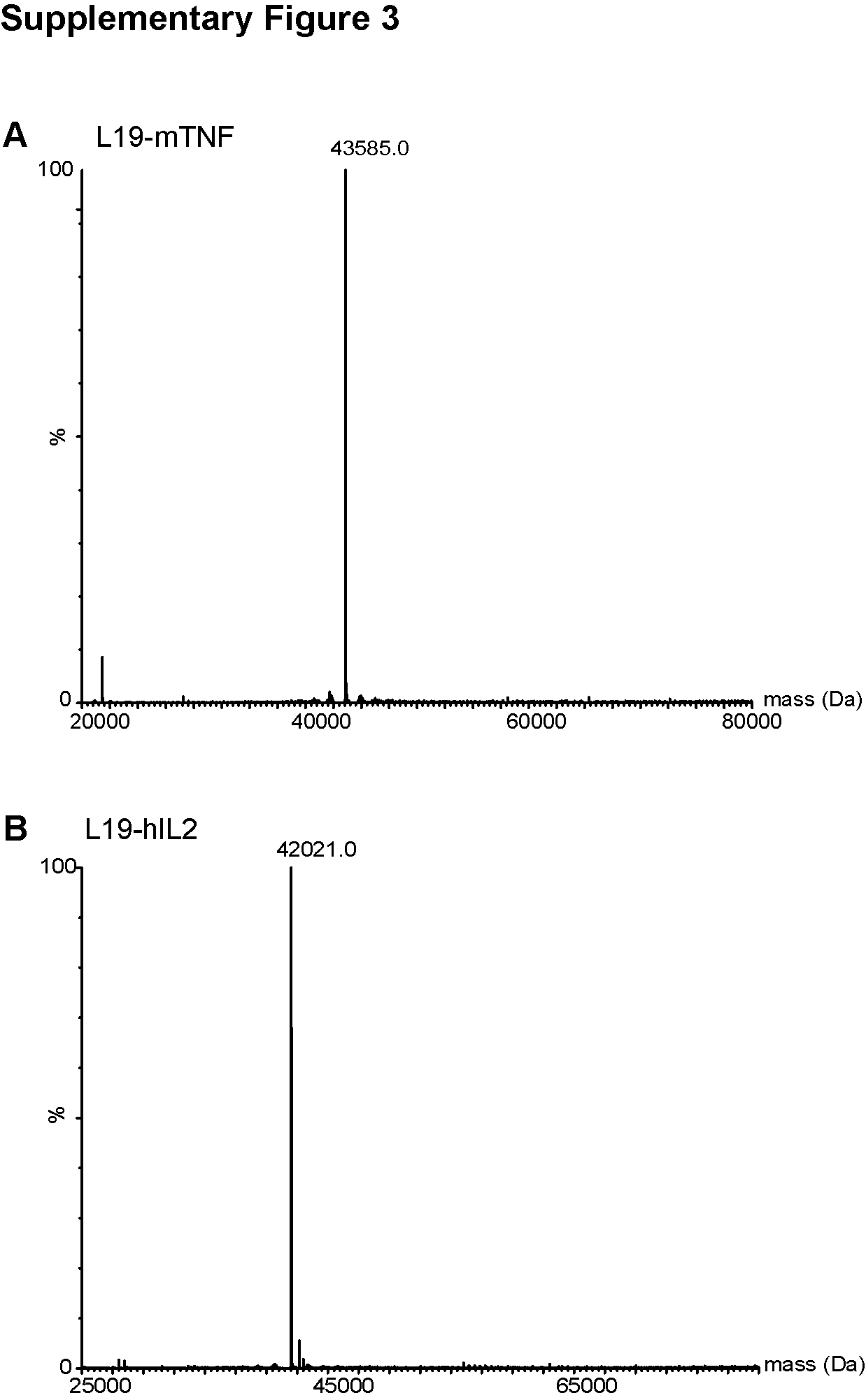

[**Supplementary Figure 3**] ESI/LC-MS of monomeric L19-mTNF [**A**] and monomeric L19-hIL2 [**B**].

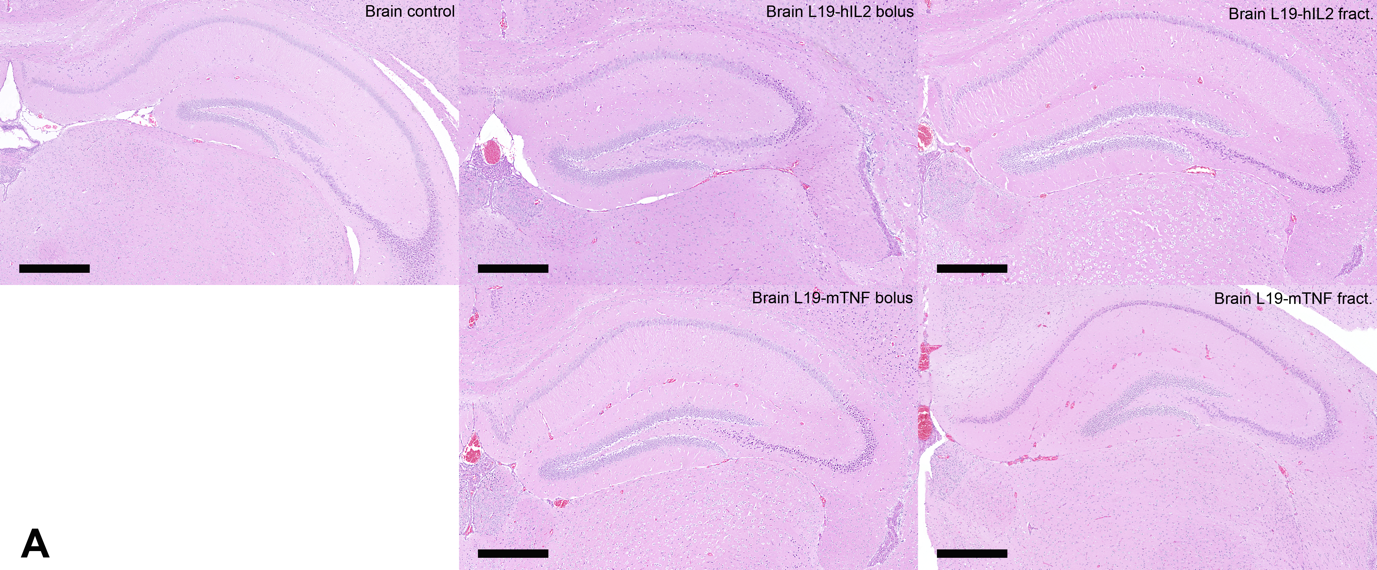

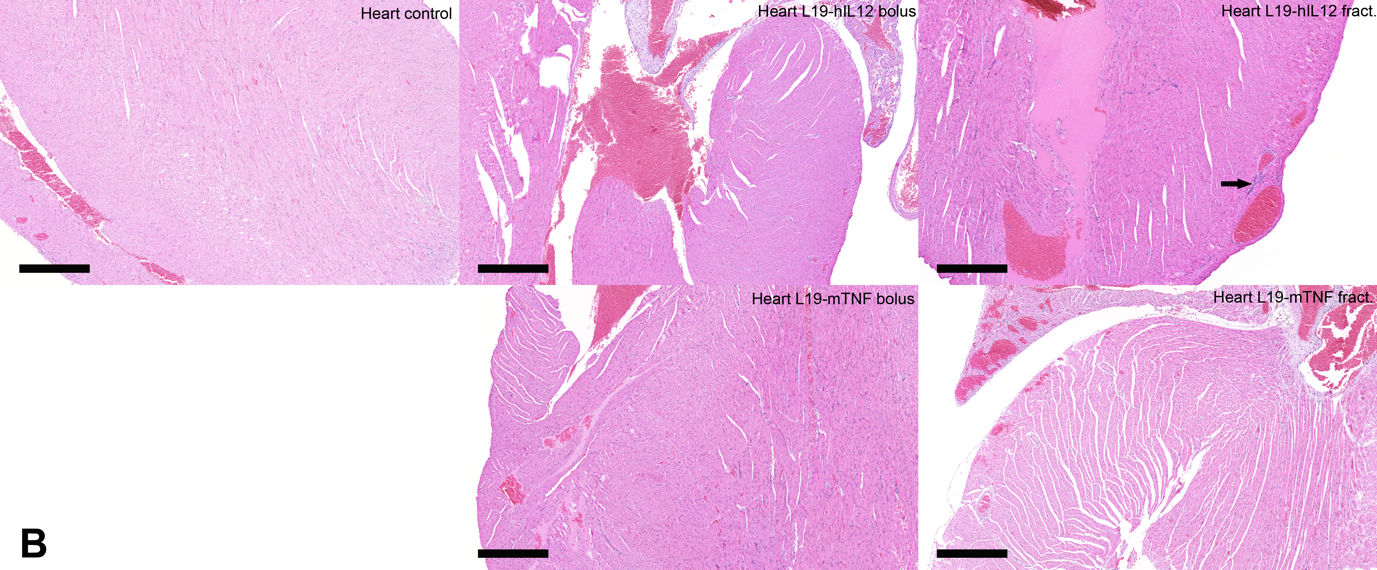

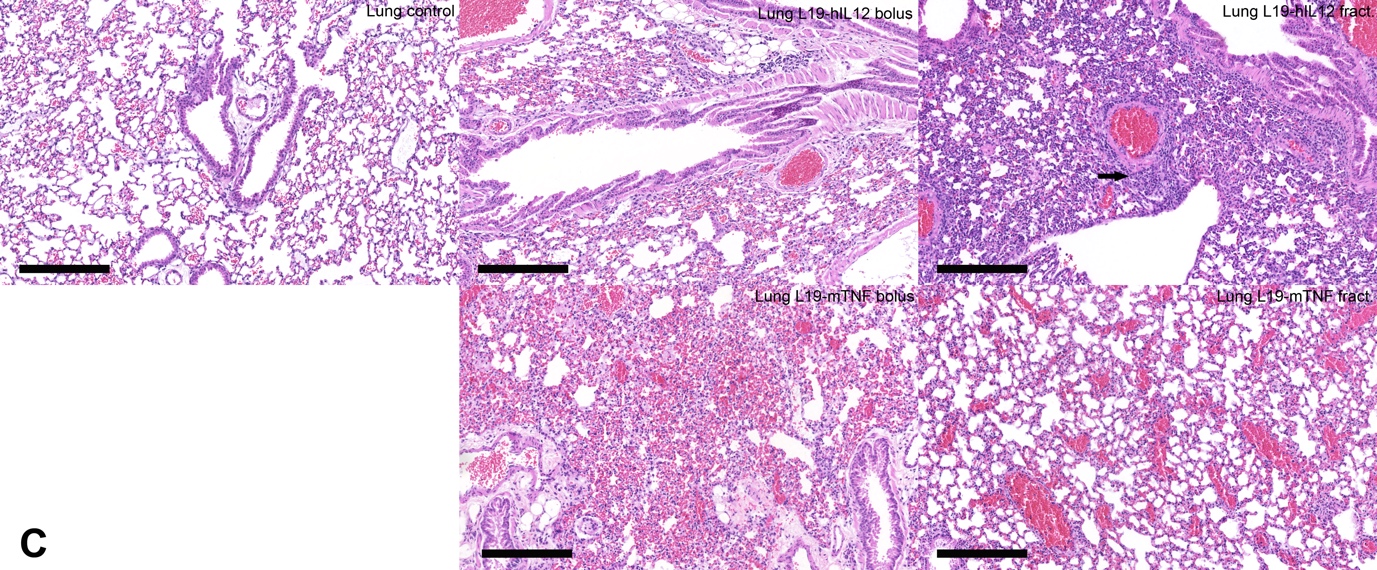

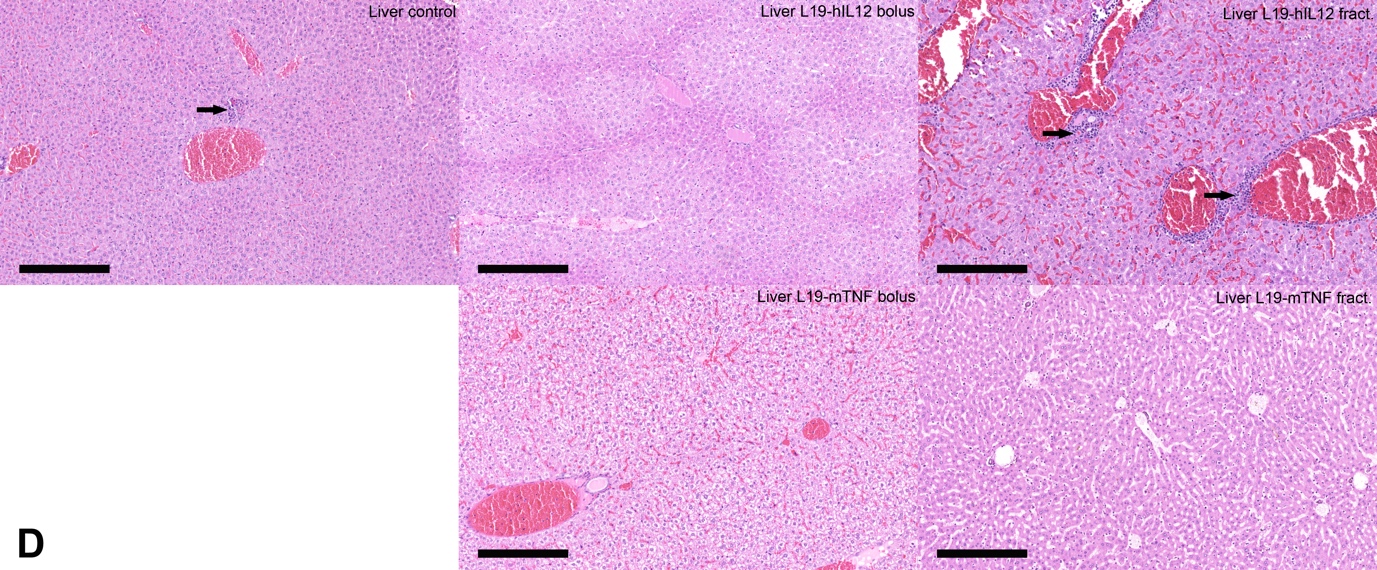

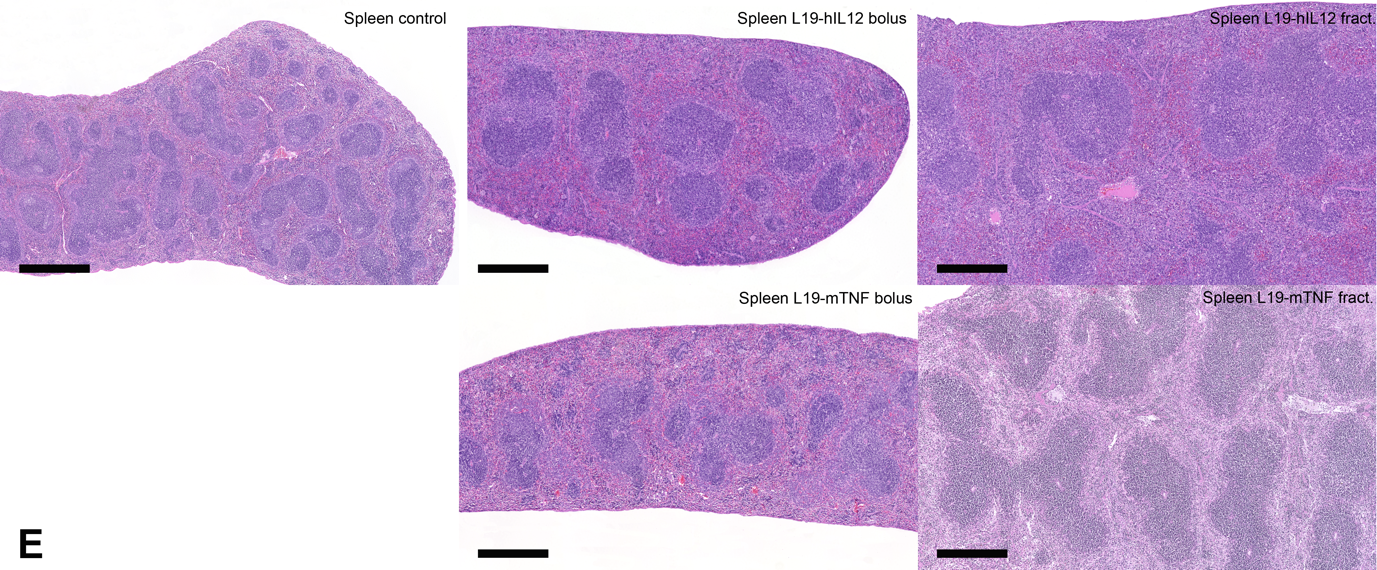

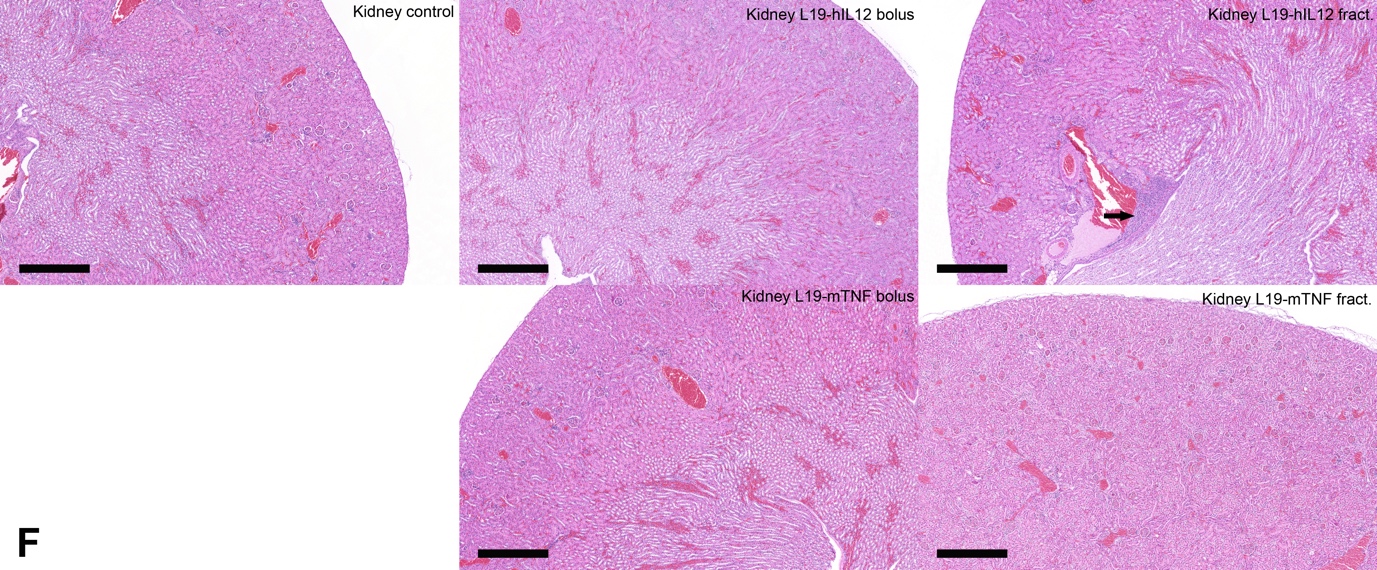

[**Supplementary Figure 4**]**:** H&E staining of organ sections of mice that received intravenous injections of saline, L19-hIL2 or L19-mTNF. Mice were sacrificed 24h after a single bolus administration of 400 µg L19-hIL2, 5 µg L19-mTNF or saline, and 24h after three fractionated injections of 200 µg L19-hIL2 or 3 µg L19-mTNF. [**A**] Brain (hippocampus) of control animal and treated mice without histopathological findings. Scale bar = 500 µm. [**B**] Heart of control animal and treated mice. Minimal lymphocytic infiltrates (arrow) in L19-hIL12 fractionated mouse. All other animals did not show histopathological findings. Scale bar = 500 µm. [**C**] Pulmonary tissue of control animal and treated mice. Mild interstitial perivascular lymphocytic infiltration (arrow) in in L19-hIL12 fractionated mouse. All other animals did not show significant histopathological findings. Scale bar = 250 µm. [**D**] Liver of control animal and treated mice. Control animal with focal necrosis (left arrow). Mild multifocal periportal lymphocytic infiltration (right arrows) in L19-hIL12 fractionated mouse. Scale bar = 250 µm. [**E**] Spleen of control animal and treated mice. All animals showed mild to moderate lymphoid hyperplasia and extramedullary hematopoiesis in the red pulp. Scale bar = 500 µm. [**F**] Kidney of control animal and treated mice. Mild interstitial lymphocytic infiltration (arrow) in the pelvic region of a L19-hIL12 fractionated mouse. All other animals did not show significant histopathological findings. Scale bar = 500 µm.
